# Pick Your Poison: Benzalkonium Chloride and Copper Enable Nanocellulose Derivatives to Form Antimicrobial Properties Against a Spectrum of Microorganisms

**DOI:** 10.1101/783076

**Authors:** Matthew J. Winans, Jennifer E.G. Gallagher, Jacek Jaczynski, Gloria Oporto

## Abstract

TEMPO nanofibrillated cellulose (TNFC), nanofibrillated cellulose (NFC), carboxymethyl cellulose (CMC), and lignin were used as templates for the addition of two well-known antimicrobial substances, benzalkonium chloride (BZK) and copper. The resulting hybrid of cellulose and antimicrobial materials were analyzed for biocidal activity in three separate products. Assays of the nanocellulose-antimicrobials were assayed for viability against *Escherichia coli* in suspension, against *Saccharomyces cerevisiae* on PVA plastic, and against *bacillus lincheniformis* in paper additives. Instant biocidal activity was achieved with a minimal inhibitory concentration of 0.116 M BZK-TNFC hybrid suspension. BZK-Lignin and BZK-CMC suspensions demonstrated increased antimicrobial activity with longer exposure times during a 24-hour exposure which completely inhibited the bacteria. BZK was slowly released into the suspension, a desirable trait for long-term antimicrobial activity. PVA plastic incorporated with BZK/Cu-nanocellulose scaffolds created solid films that completely inhibited yeast growth by 270 seconds. Interestingly, lignin-BZK PVA films counteracted each other and showed no biocidal activity at all. The multiple combinations of nanocellulose and biocidal agents in the surface viability assay demonstrates the importance of synergy between both components in designing nanocellulose antimicrobials. TNFC-Cu was more suited to inhibit growth in paper than NFC-Cu as seen in a zone of inhibition assay. The most potent biocidal material in PVA was NFC-BZK. Here we show the diversity of the cellulosic derivatives and their impact on the antimicrobial additive. We employed a variety of assays to assess to biocidal of these nanoparticles against three species of bacteria and yeast relevant to food packaging and medical fields. From our study, there are many factors that play a role in the design of antimicrobial materials; cellulose derivative scaffold, antimicrobial agent, type of final material in which to be incorporated, target organism, and duration of application.

## INTRODUCTION

Cellulose has an extensive range in industrial and technological applications that includes the production of paper, household commodities, and development of advanced biomedical tissue engineering scaffolding (Haroun *et. al* 2009; Yang *et. al* 2011). The natural abundance, innate nontoxicity, and biodegradability of cellulose makes it a desirable material, but its lack of antimicrobial properties limit their spectrum of application. Many research groups have focused on improving this specific shortcoming by affixing additional molecules that exhibit antimicrobial behavior. Cellulose is a suitable template for nanomaterials because of its large number of functional groups available for chemical modification (Cai *et. al* 2009; Shin *et. al* 2007). Nanocellulose production has primarily centered on the incorporation of nonorganic metallic nanoparticles (Diez *et. al* 2011; Silva *et. al* 2014; Zhong *et al.* 2013). The untapped potential of organic compounds such as benzalkonium chloride (BZK) and copper remains unexplored in nanocellulose science and may prove appropriate for biomedical applications.

Benzalkonium chloride (BZK) is an organic salt that is well known for its biocidal activity against bacteria, fungi, protozoa, and some viruses. BZK is positively charged and acts as a cationic surfactant interacting with the negatively charged cell wall and the plasma membranes. This interaction permits leakage of small molecules, degradation of biomolecules, and eventual lysis facilitating cell death (Krishnan et. al., 2018). BZK is stable in a wide range of conditions and is commonly used as a bioactive agent in many pharmaceutical, personal care, and disinfectant consumer products. Unlike many other organic molecules with similar biocidal properties, BZK’s toxicity to humans is low and allows for topical medicinal application (Thorsteinsson *et. al* 2003). BZKs effectiveness in the synthesis of antimicrobial cellulose biocomposites has been established in previous research. The incorporation of a chitosan-BZK complex into microfibrillated cellulose increased strength and antimicrobial properties of sodium alginate films, commonly utilized for food packaging applications (Liu *et. al* 2013). Additionally, through the immersion of freeze-dried bacterial cellulose films in a BZK solution, the resulting cellulose-BZK hybrid films effectively inhibited *Staphylococcus aureus* and *Bacillus subtilus* for 24 hours at low concentrations (Wei *et. al* 2011).

This research took a novel approach to the hybridization of cellulose and BZK by utilizing TEMPO (2,2,6,6-tetramethylpiperidine-1-oxyl)-oxidized nanofibrillated cellulose (TNFC) and carboxymethyl cellulose (CMC) to entrap BZK molecules in a hybrid suspension. TNFC and CMC are distinguished from other cellulose derivatives by their high surface area ratio and the presence of converted carboxyl groups that aid in the physical binding of organic molecules (Isogai *et. al* 2011). The antimicrobial properties of the resulting suspensions were tested against a gram-negative bacteria, *Escherichia coli*. Furthermore, these cellulose scaffolds were complexed with copper as a paper/fiber additive and were tested against gram-positive bacteria, *bacillus lincheniformis*, as a representative for nosocomial infections and biofilm formation in biomedical applications. Lastly, antimicrobial properties of the nanocellulose-Cu/BZK embedded in polyvinyl alcohol (PVA) films were assayed through surface viability against *Saccharomyces cerevisiae* as a representative of spoilage in packaged foods (Jiang, *et. al* 2016). Solid cellulose-BZK film material was obtained through a solvent exchange and freeze-drying procedure, previously detailed (Zhong *et. al* 2015). Films were characterized using inverse gas chromatography (IGC). PVA film synthesis and viability assessment demonstrated the feasibility of nanocellulose-antimicrobial complexes in food packaging and medical applications by providing microbial biocidal activity through a controlled release system.

## RESULTS AND DISCUSSION

### TNFC-BZK hybrid suspension antimicrobial properties

*E. coli* was exposed to 0.116 M TNFC-BZK hybrid suspension, washed, and plated on normal growth media to assess the antimicrobial activity. The concentration of *E. coli* in suspension prior to dilution was 0.8 × 10^7^ CFU/mL. *E. coli* was serially diluted to 10^−7^ prior to spread plating from treatment with TNFC and pristine PBS (Figure 1A). There were 8 visible colonies in the untreated (pristine PBS) and 8 visible colonies in the TNFC treated solution. *E. coli* treated with TNFC-BZK was serially diluted to 10^−2^ prior to spread plating and no visible colonies were formed, indicating the complete inhibition of *E. coli*. Given the identical amounts of TNFC in both treatments, the antimicrobial action can be attributed to BZK.

**Figure 1.**
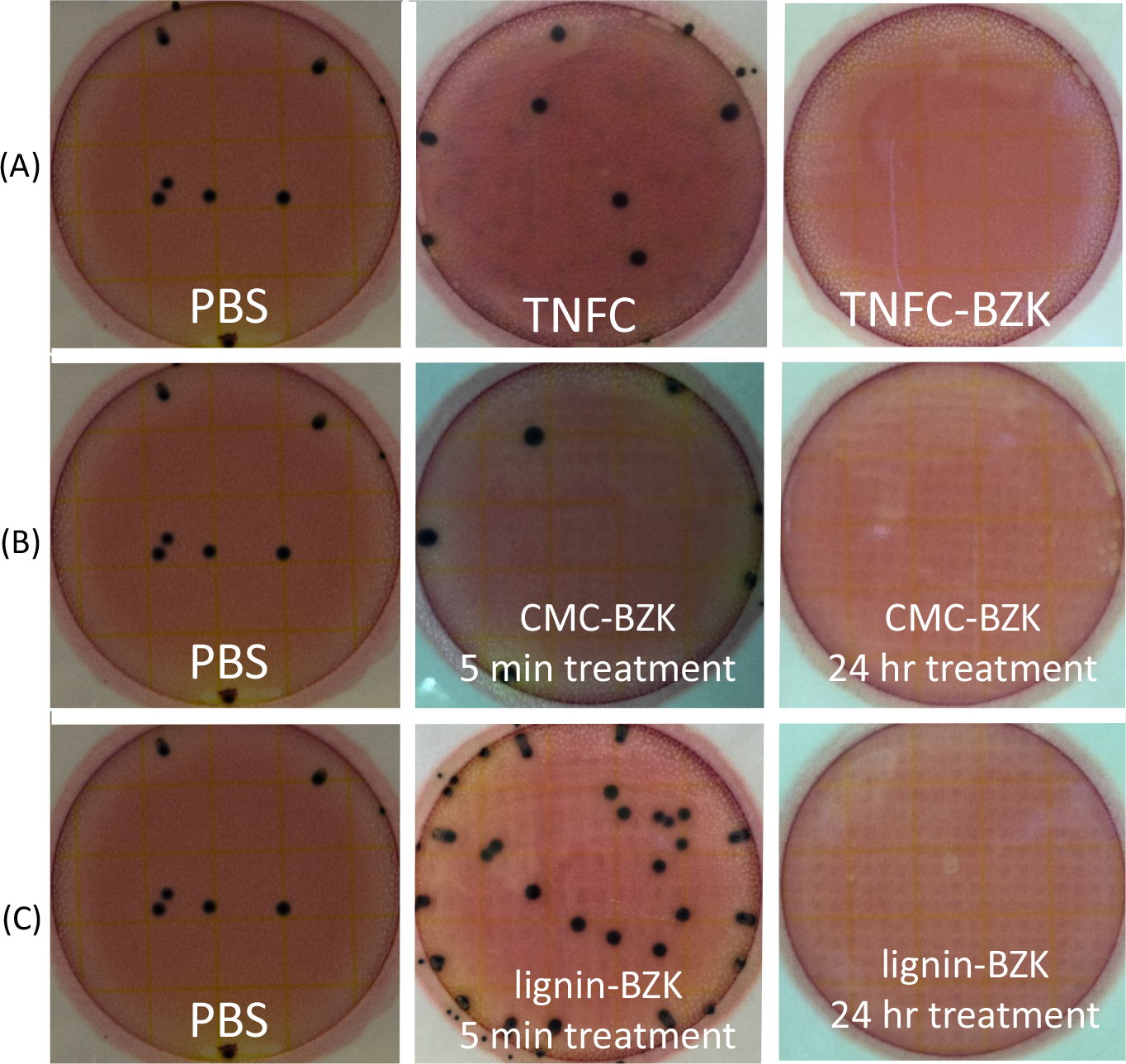
*E. coli* colony forming units assessment. (A) Enumeration of *E. coli* treated with PBS, TNFC, and TNFC-BZK hybrid suspensions on 3M *E. coli*/Coliform Count Petrifilm. *E. coli* was serially diluted for spread plating from 0.8 × 10^7^ CFU/mL to 10^−7^ for PBS and TNFC cultures. *E. coli* was serially diluted for spread plating from 0.8 × 10^7^ CFU/mL to 10^−2^ for TNFC-BZK cultures. (B) Enumeration of *E. coli* treated with PBS, CMC-BZK after 5 mins, and CMC-BZK after 24 hrs. on 3M *E. coli*/Coliform Count Petrifilm. *E. coli* was serially diluted for spread plating from 0.8 × 10^7^ CFU/mL to 10^−6^ for CMC-BZK cultures. (C) Enumeration of *E. coli* treated with PBS, lignin-BZK for 5 mins, and lignin-BZK for 24 hrs on 3M *E. coli*/Coliform Count Petrifilm. *E. coli* was serially diluted for spread plating from 0.8 × 10^7^ CFU/mL to 10^−6^ for lignin-BZK cultures.

### CMC-BZK and Lignin-BZK hybrid suspension antimicrobial properties

To assess the role of different cellulose scaffolds on antimicrobial effectiveness and timeliness, CMC and lignin were used instead of TNFC. Enumeration of *E. coli* treated with pristine PBS, 0.046 M CMC-BZK suspension for 5-minutes, and 0.046 M CMC-BZK hybrid suspension for 24-hours was conducted (Figure 1B). *E. coli* was serially diluted to 10^−7^ in preparation for spread plating PBS treatment. *E. coli* was serially diluted to 10^−6^ in preparation for spread plating the CMC-BZK timed treatments. When *E. coli* was treated for 5 min with 0.046 M CMC-BZK, 0.6 × 10^6^ CFU/mL was present after treatment. There were no visible colonies after 24 hours, indicating the complete inhibition of *E. coli* at this concentration.

To assess the role of different cellulose scaffolds on antimicrobial effectiveness and timeliness, Lignin-BZK suspensions were used instead of TNFC. Enumeration of *E. coli* treated with pristine PBS, 0.046 M Lignin-BZK suspension for a 5 minutes, and Lignin-BZK suspension for 24 hours was conducted (Figure 1C). *E. coli* was serially diluted to 10^−7^ in preparation for spread plating PBS treatment. *E. coli* was serially diluted to 10^−6^ in preparation for spread plating the lignin-BZK timed treatments. When the *E. coli* was treated for 5 minutes with lignin-BZK hybrid suspension, 3.1 × 10^6^ CFU/mL were detected, while there were no visible colonies at 10^−6^ for CMC-BZK cultures after 24-hrs of treatment. These results, in combination with results from CMC testing, indicate an increase in antimicrobial properties associated with longer exposure times.

### Quantification of surface viability

To assess the antimicrobial abilities of the nanocellulose-biocidal complexes in PVA plastic against yeast, changes in *S. cerevisiae* viability were recorded. The materials were incorporated into polyvinyl alcohol (PVA) films using a solvent casting method. *S. cerevisiae* RM11, a hardy agricultural isolate, was exposed to the final films. NFC films, complexed with either biocidal agent, were the most potent material with no viable yeast after 30 seconds of exposure while 90 seconds of exposure was required for NFC-Cu (Figure 2). Carboxymethyl cellulose was the least effective cellulose scaffold for acute exposure and second least effective in the overall BZK exposure trials. Lignin-Cu material provided substantial decrease in viability. For BZK additions, lignin resulted in the least effective hybrid material with no effect on viability. The lignin-BZK film caused no cellular death, completely countering the effect of BZK as an antimicrobial agent. More research is needed to determine the rescuing effect observed in this study. TNFC-BZK closely mirrored the highly toxic rate of cellular death of NFC-BZK. When TNFC is coupled with copper, it does not mirror the high toxicity that NFC-Cu does. The difference seen is suggested to be from the synergistic interaction between cellulose scaffold and the biocidal agent.

**Figure 2.**
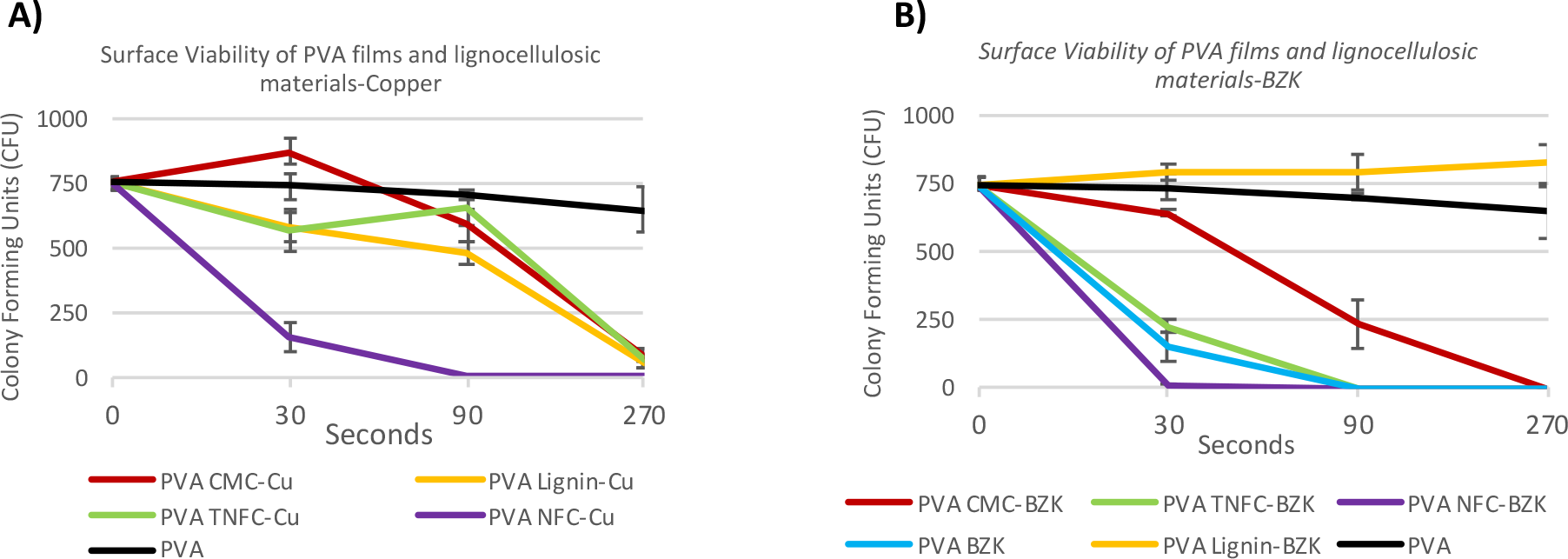
Viability of *S. cerevisiae* yeast in contact with PVA surfaces embedded with nanocellulose-biocidal materials. Colony forming units were assayed for each surface material in the indicted exposure time in seconds.

The difference in synergy between scaffold and biocidal agent suggests that pairing of the scaffold and agent is an important consideration for its intended use as a PVA plastic. Copper generates free radicals through Fenton reactions when it interacts with water and previous work demonstrated the antimicrobial activity of CMC-Cu hybrid suspensions against *S. cerevisiae* (Rong-Mullins *et al.* 2017). BZK disrupts the cellular membrane’s surface charge and bacterial studies have shown that *Pseudomonas aeruginosa* copes with benzalkonium chlorides (BAC) in three ways; decreasing porin expression related to BAC, increasing efflux pump activity, and stabilizing the cell surface charge (Krishnan *et al.* 2018). Thus, these biocidal agents have different modes of action that require consideration when designing antimicrobial cellulose materials.

### Zone of inhibition

To assess the antimicrobial abilities of these materials incorporated into paper products, a zone of inhibition assay was conducted. *Bacillus licheniformis is* a bacteria responsible for a range of clinical conditions and food spoilages (Salkinoja-Salonen *et al.* 1999). TNFC-Cu paper discs created the largest zone of inhibition at 2.72 cm with a standard deviation of 0.11 followed by NFC-Cu at 2.01 cm with a standard deviation of 0.15 (Figure 3). A student’s T test resulted in p-value of 1.1 × 10^−17^. CMC-Cu discs did not show any zones of inhibition, but several discs showed what appeared to be an immediate ring of slightly retarded growth at an average diameter of 1.7 cm with a standard deviation of 0.16 (results not shown). The CMC-Cu growth was distinctly different from the inhibition displayed by TNFC-Cu and NFC-Cu materials. The zone of inhibitions of each cellulose derivative was markedly different, which reflects the importance of the variation in cellulose scaffolds while constructing antimicrobial materials.

**Figure 3.**
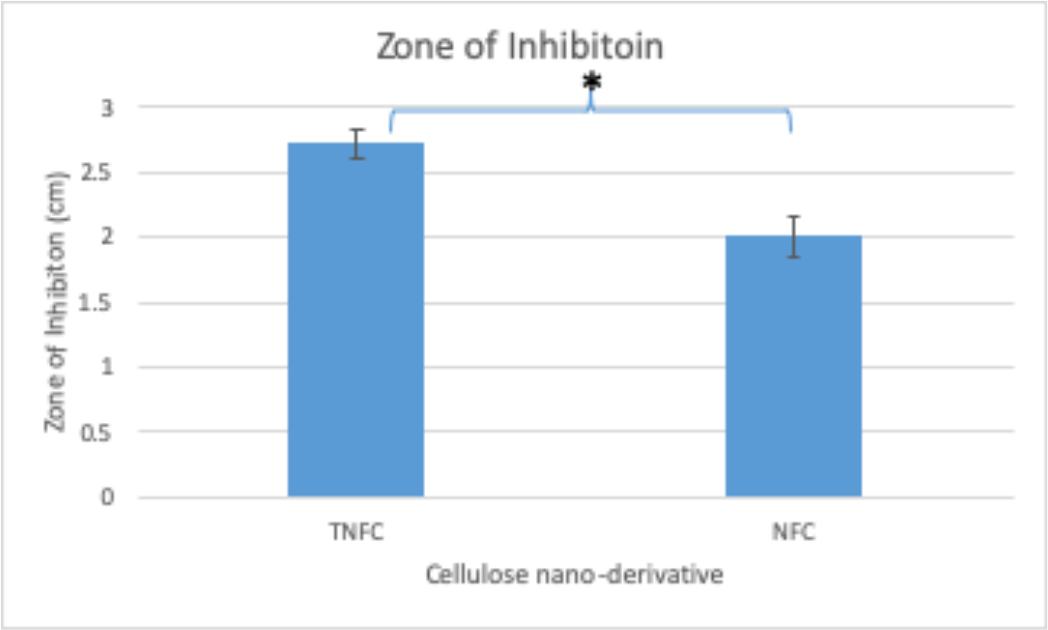
Zone of inhibition for *B. lincheniformis* against cellulose derivatives and copper hybrid materials infused in paper. In the disc diffusion assay, larger zones of inhibition represent increased toxicity of the substance against *B. lincheniformis.* Zone of inhibition measured by their diameter in ImageJ. Students T test performed in Excel, p= 1.1 × 10^−17^. Bacillus lincheniformis (ATCC 14580) on Nutrient Broth after 2 days growth.

## CONCLUSIONS

BZK was successfully used as an organic biocide to improve the antimicrobial properties of modified cellulose against bacteria and fungi. The suspensions of CMC-BZK or Lignin-BZK at 0.046 M sufficiently inhibited nonpathogenic *E. coli* after 24 hours of exposure. Here, we presented evidence of a controlled release mechanism for BZK particles in suspension. The increased antimicrobial properties seen by increasing exposure times supports a controlled release system. BZK complexed with NFC as the scaffold improved the ability to inhibit fungi compared to BZK alone in our surface viability assay. Both BZK and copper result in the eventual lysis of fungi, but through different mechanisms. Inside yeast, the copper released from CMC-Cu induces reactive oxygen species (ROS) causing free radical damage to biological molecules including protein, DNA, and lipids (Rong-Mullins, *et al.* 2017). BZK is an amphiphilic molecule and interaction with cellular membranes induces membrane disorganization, leakage of intracellular material, and degradation of nucleic acids/proteins leading to cell death. Copper is an important fungicide and through repeated exposure many agricultural yeast strains are copper tolerant (Rong-Mullins, *et al.* 2017). The most common mechanism of copper tolerance is a genomic amplification of the *CUP1* locus (Karin et al. 1984). The *CUP1* gene encodes a metallothionein that binds excess copper, preventing the production of free radicals (Karin et al. 1984). It is unknown if yeast can also become BZK resistant. In bacteria, mutations that reduce the plasma membrane’s negative charge and increase efflux pumps increase resistance during long exposures to sublethal levels (Krishnan *et al.* 2018). The nearly immediate inhibition of all yeast to the NFC-BZK films reduce the risk of selecting resistant pathogens during normal use. Hybrid suspension added to PVA films would increase the value and antimicrobial properties for food packaging applications. From our study, there are many factors that play a role in the design of antimicrobial materials; cellulose derivative scaffold, antimicrobial agent, type of final material in which to be incorporated, target organism, and duration of application.

## METHODS & MATERIALS

### Materials

TNFC gel (0.96 wt. % TNFC) was supplied by the Forest Product Laboratory, Madison, WI. PVA (99–100 % hydrolyzed, approximate molecule weight 86000) obtained from Acros Organics, USA. Sterile trypticase soy broth (TSB) was purchased from Becton–Dickinson, USA. *E. coli*/Coliform Count Petrifilm Plates obtained from 3M, USA. Butterfield phosphate buffer obtained from Hardy Diagnostics, USA. *Escherichia coli* DH5α (hereafter called *E. coli*) was purchased from Invitrogen Inc., USA. Sodium carboxymethyl cellulose (CMC) (average molecular weight: 90,000) obtained from Sigma Aldrich, USA. Ethanol (reagent alcohol, specially denatured alcohol formula 3A 95 % and Isopropyl alcohol 5 % by volume) obtained from Sigma Aldrich, USA. Tert-butanol (99.5 %, for analysis) obtained from Acros Organic, USA. Phosphate buffer solution (PBS) obtained from Hardy Diagnostics, Santa Maria, CA, USA. Deionized water was used throughout the study. *S. cerevisiae* strain, RM11, is derived from an agricultural isolate (Mortimer and Polsinelli 1999). Yeast were grown in YPD media (1% yeast extract, 2% peptone, 2% dextrose). Yeast extract and peptone obtained from DB Difco. Dextrose obtained from Fischer Scientific.

### Methods

#### Preparation of hybrid material and hybrid PVA films

TNFC gel (15 mL) was dissolved in deionized water (35 mL). The hybrid suspension was prepared by the addition of 10 mL BZK solution (25% w/w) to the TNFC solution under vigorous stirring. The suspension was stirred for an additional two hours to facilitate the dispersion and binding of BZK to the cellulosic backbone.

PVA films were cast by preheating deionized water (90 mL) to dissolve solid PVA particles (10 g). The TNFC-BZK hybrid suspension was added dropwise to the PVA solution under vigorous stirring and the resulting mixture was stirred for an additional 2 hours. The PVA was then cast in thin layers into Petri dishes and degassed in a desiccator for 24 hours. After degassing, the films were allowed to rest at room temperature for 12 hours before being transferred to a dry oven (50° C) for 24 hours.

For CMC and Lignin hybrids, solid cellulosic material was added to deionized water (50 mL) and stirred until fully dissolved. A lower concentration of BZK (10% w/w) was added to both solutions due to the limited availability of CMC to complex with BZK. All other hybrid suspensions and films were prepared with the same procedure used for TNFC suspensions detailed above.

#### Antimicrobial testing of hybrid suspensions

*E. coli* was cultured in TSB media for 24 hrs. at 37°C with agitation set to 150 rpm. 1 mL aliquots of the saturated culture were used for antimicrobial testing. 1 mL of saturated *E. coli* culture was treated by addition to 9 mL liquid suspension, thoroughly mixed, and then treated for 5 min. Enumeration occurred after a ten-fold serial dilution and spread plating (Black et. al 2006). 1 mL of the *E. coli*/hybrid suspension was added to 9 mL PBS and homogenized. Subsequent serial dilutions were performed in PBS. 1 mL aliquots of each dilution were enumerated using 3M Petrifilm *E. coli* counting plates after incubation at 35°C for 48 hours.

#### Surface viability testing of PVA-BZK hybrid films

The BZK-cellulose PVA films were tested for antimicrobial properties against the budding yeast, *Saccharomyces cerevisiae* strain RM11, using a surface viability protocol. Saturated overnight cultures of yeast were spun at 3000 rpm for 3 minutes, washed in phosphate-buffered saline (PBS) and resuspended in PBS solution. Optical density_600nm_ readings were taken using a Thermo Spectronic model GENESYS 10 UV. Using separated pipette tips, 29,000 *S. cerevisiae* cells in 20 μl were deposited on the surface and spread across the 1 cm^2^ PVA film surface. PVA films were prewashed in ethanol and air-dried in a chemical fume hood. Data was collected for exposure times of 5, 60, 120, and 300 seconds. Additionally, the untreated control was labeled as timepoint 0. After the designated exposure times, 10ml of PBS was used to wash the coupon and harvest all cells into a 50 ml centrifuge tube. The resultant solution with coupon was vortexed and the coupon was discarded. 50 μl of the treated cells/PBS suspensions were plated onto YPD. Each plate was incubated at 30°C for 48 hr. and then colony forming units (CFU) were counted manually by eye.

#### Zone of inhibition

Cellulose-copper materials were embedded via filter paper application and allowed to air dry. A hole punch, producing ¼ inch diameter discs, was used. Nutrient broth (0.1% beef broth, 0.2% yeast extract, 0.5% peptone, 0.5%NaCl) was used to culture *Bacillus lincheniformis* (ATCC 14580). Overnight cultures were grown to saturation and diluted by optical density_600nm_ to 0.5 and aseptically spread onto nutrient broth plates. Once dried, the punch-out discs were then placed in triplicate on each inoculated plate. The plate was cultured for 2 days at 37°C. Pictures were taken after growth had occurred. Pictures were analyzed with ImageJ and each disc’s zone of inhibition was measured twice, totaling 6 measurements per treatment. The average and the standard deviation were calculated.

## ACKNOWLEDGEMENTS

Funds for this project were provided by USDA-NIFA Grant No. 2013-34638-21481 “Development of novel hybrid cellulose nanocomposite film with potent biocide properties utilizing low quality Appalachian hardwoods”. MJW was supported by The National Science Foundation’s grant “IGERT: REN@WVU - Research and Education in Nanotoxicology at West Virginia University”, award number 1144676. MJW conducted yeast surface viability and zone of inhibition experiments. Oliver Lin assisted in the viability studies on yeast. JJ & GO conducted *E. coli* studies.

